# Translating GWAS Findings Into Therapies For Depression And Anxiety Disorders: Drug Repositioning Using Gene-Set Analyses Reveals Enrichment Of Psychiatric Drug Classes

**DOI:** 10.1101/132563

**Authors:** Hon-Cheong So, Alexandria Lau, Carlos Kwan-Long Chau, Sze-Yung Wong

## Abstract

Depression and anxiety disorders are the first and sixth leading cause of disability worldwide according to latest reports from the World Health Organization. Despite their high prevalence and the significant disability resulted, there are limited advances in new drug development. On the other hand, the advent of genome-wide association studies (GWAS) has greatly improved our understanding of the genetic basis underlying psychiatric disorders.

In this work we employed gene-set analyses of GWAS summary statistics for drug repositioning. We explored five related GWAS datasets, including two on major depressive disorder (MDD-PGC and MDD-CONVERGE, with the latter focusing on severe melancholic depression), one on anxiety disorders, and two on depressive symptoms and neuroticism in the population. We extracted gene-sets associated with each drug from DSigDB and examined their association with each GWAS phenotype. We also performed repositioning analyses on meta-analyzed GWAS data, integrating evidence from all related phenotypes.

Importantly, we showed that the repositioning hits are generally enriched for known psychiatric medications or those considered in clinical trials, except for MDD-PGC. Enrichment was seen for antidepressants and anxiolytics but also for antipsychotics. We also revealed new candidates or drug classes for repositioning, some of which were supported by experimental or clinical studies. For example, the top repositioning hit using meta-analyzed p-values was fendiline, which was shown to produce antidepressant-like effects in mouse models by inhibition of acid sphingomyelinase and reducing ceramide levels. Taken together, our findings suggest that human genomic data such as GWAS are useful in guiding drug discoveries for depression and anxiety disorders.

## Introduction

Depression and anxiety disorders are among the most common psychiatric disorders. According to the latest report by the World Health organization, depression affects more than 300 million people worldwide and is the leading cause of disability ^1^. Anxiety disorders affects more than 260 million people and is the sixth leading cause of disability ^1^. The two disorders are highly comorbid and might share common pathophysiologies^2,3^. Nevertheless, pharmacological treatment for major depressive disorder (MDD) or anxiety disorders (AD) has not seen much advance in the last two decades or so, with a lack of therapies having novel mechanisms of action. In addition, only about one third of MDD patients achieve complete remission after a single antidepressant trial ^4^ and around 10 to 30% of patients are treatment-resistant ^5^.

On the other hand, with the advent of high-throughput technologies such as genome-wide association studies (GWAS) in the last decade, we have gained a much better understanding of the genetic bases of many complex diseases. It is hoped that human genomics data will accelerate drug development for psychiatric disorders, especially due to the difficulties for animal models to fully mimic human psychiatric conditions such as depression ^6^.

We hypothesize that gene-sets associated with antidepressants or anxiolytics, or more generally with psychiatric medications, will be enriched among the GWAS results of depression and anxiety phenotypes. If the gene-set analysis (GSA) approach is able to “re-discover” known treatments, it might also be able to reveal new therapies for depression and anxiety disorders. Practically, if the pathway targeted by a particular drug (or chemical) is significantly enriched in GWAS but the drug is *not* a known treatment for the disease, it may serve as a good candidate for repositioning.

With regards to GWAS-based drug discovery, the current focus is mainly on identification of new drug targets from the top GWAS hits. In an earlier study, Sanseau et al. ^7^ identified the most significant GWAS hits from a range of diseases and compared them against known drug targets to find “mismatches” (i.e. drug indication different from the studied disorder) as candidates for repurposing. While it is intuitive and legitimate to focus on the most significant SNPs, for many complex traits the genetic architecture may be highly polygenic and variants of weaker effects may be “hidden”. Moreover, given the complex and multifactorial etiologies of many complex diseases, the development of multi-target drugs with wide-ranging biological activities (known as “polypharmacology”) is gaining increased attention (please refer to e.g. Anighoro et al. ^8^ for a review). It is argued that multi-target drugs may have improved efficacy over highly selective pharmacological agents, as they tackle multiple pathogenic pathways in the system. A gene-set or pathway based approach to drug repositioning follows this multi-target paradigm.

Gene-set analysis is an established approach to gain biological insight into expression microarrays, GWAS or other high-throughput “omics” studies ^9,10^. De Jong et al. made use of GSA to identify repurposing opportunities for schizophrenia ^11^ and several top results were supported by the literature. In another very recent study, Gasper et al. ^12^ performed further analyses of GSA results, and reported that GWAS signals of schizophrenia are enriched for neuropsychiatric medications as sample size increases. Another related study on schizophrenia was conducted by Ruderfer et al. ^13^, who collected genome-wide significant GWAS variants and exome sequencing results and compared the identified genes against known or predicted drug targets. Significant enrichment was observed for antipsychotics.

In this study we take a different focus on depression and anxiety disorders, which are highly prevalent and disabling disorders. Besides considering individual GWAS studies of phenotypes related to depression and anxiety, we also performed analyses on combined GWAS summary statistics to improve the power.

## Methods

### Overview of analytic approach

Our analyses can be broadly divided into two steps. Firstly, we extracted gene-sets associated with a variety of drugs (or chemicals), and tested whether the gene-set associated with *each individual drug* is enriched among the GWAS results. The drugs ranked among the top (i.e. those with lower *p*-values) were considered potential candidates for repositioning. In the second step, we performed analyses on the drugs. We evaluated the prioritized drugs and tested which *drug classes* were enriched. As we have hypothesized above, we would specifically test whether antidepressants/anxiolytics and other psychiatric medications were enriched among the repositioning results.

### Genome-wide association studies data

We considered five GWAS datasets that are associated with depression and anxiety. Two are studies of major depressive disorder (MDD), namely MDD-PGC ^14^ and MDD-CONVERGE ^15^. However, the two studies are different in a number of ways. The MDD-PGC sample (*N* = 18,759) is composed of Caucasians of both sexes, while MDD-CONVERGE (*N* = 10,640) is a cohort of Chinese women. The MDD-CONVERGE sample mainly consists of hospital-ascertained cases affected by severe depression, of whom ~85% had melancholic symptoms ^15^. The MDD-PGC sample on the other hand is more heterogeneous and not specifically enriched for any subtypes of depression ^14^

Another two GWAS studies were meta-analyses on depressive symptoms and neuroticism conducted by the Social Science Genetics Association Consortium (SSGAC) ^16^. The meta-analysis on depressive symptoms (SSGAC-DS) included the MDD-PGC study (*N* = 18,759) and a case-control sample from the Genetic Epidemiology Research on Aging (GERA) Cohort (*N* = 56,368), but it also comprised an UK BioBank sample made up of general population (*N* = 105,739). Depressive symptoms were measured by a self-reported questionnaire. We also included another study on neuroticism (SSGAC-NEU) (*N* = 170,906), as this personality trait is known to be closely associated with depression and anxiety disorders ^17^. In addition, antidepressants may affect personality traits, including a reduction in neuroticism, independent of their effects on depressive symptoms ^18^.

The fifth dataset is a GWAS meta-analysis of anxiety disorders, including generalized anxiety disorder, panic disorder, social phobia, agoraphobia, and specific phobias ^19^. We extracted the GWAS results of the case-control analyses (*N* = 17,310).

GWAS summary results were downloaded from https://www.med.unc.edu/pgc/results-and-downloads and https://www.thessgac.org/data.

### Extracting gene-sets associated with each drug

We made use of the DSigDB database ^20^ to extract gene-sets related to each drug. DSigDB holds gene-sets for a total of 17839 unique compounds. The gene-sets were compiled according to multiple sources: (1) bioassay results from PubChem ^21^ and ChEMBL ^22^; (2) kinase profiling assay from the literature and two kinase databases (Medical Research Council Kinase Inhibitor database and Harvard Medical School Library of Integrated Network-based Cellular Signatures database); (3) differentially expressed genes after drug treatment (with >2 fold-change compared to controls), as derived from the Connectivity Map ^23^; and (4) manually curated and text mined drug targets from the Therapeutics Targets Database (Qin et al., 2014) and the Comparative Toxicogenomics Database ^24^. We downloaded the entire database from http://tanlab.ucdenver.edu/DSigDB. The above sources captured different aspects of drug properties, for example differentially expressed genes from perturbation experiments might be different from drug target genes defined in bioassay studies. Besides performing analyses based on the whole database which incorporates a broad definition of drug-related genes, we also performed a separate enrichment analysis for genes derived from PubChem and ChEMBL only, as they represent conventionally defined “drug targets” that are more well-studied and perhaps more directly associated with drug actions.

It should be noted that although the focus is on drug “repositioning”, the analytic framework is general and can apply to any drugs with some known associated genes. Indeed DSigDB contains a substantial number of drugs which do not have an approved indication yet, which were still included in our analyses.

### Gene-set analysis (GSA) approach

We first converted the SNP-based test results to gene-based test results. We employed fastBAT^25^ (included in the software package GCTA) for gene-based analyses. FastBAT computes the sum of chi-square statistics over all SNPs within a gene and uses an analytic approach to compute the *p*-value. Gene size and linkage disequilibrium patterns are taken into account when computing the *p*-values. The same statistical approach for gene-based tests is also used by two other popular programs, VEGAS ^26^ and PLINK ^27^, although they computed *p*-values by simulations or permutations. FastBAT has been shown to be equivalent to VEGAS and PLINK at higher *p*-values (>1E-06) and more accurate than these two programs for smaller *p*^25^. We ran fastBAT with the default settings and used the 1000 Genome genotype data as the reference panel.

We then performed a standard GSA by comparing gene-based test statistics within and outside the gene-set. We adopted the same approach as implemented in MAGMA ^28^, which is also reviewed in de Leeuw et al. ^10^. Briefly, gene-based *p*-values are first converted to z-statistics by z = Φ^−1^ (*p*), where Φ^−1^ is the probit function (more negative z-values represent stronger statistical associations). We then employed a single-sided two-sample t-test to see if the mean *z*-statistics of genes within the gene-set is lower than that outside the gene-set. To avoid results driven by only a few genes, we only considered drugs with at least 5 genes in their gene-sets. A total of 5232 drugs were included for final analyses.

### Combining p-values across datasets

Besides analyzing each GWAS dataset in turn for repositioning opportunities, we also considered the aggregate contribution of all datasets, as depression, anxiety and neuroticism are closely connected to each other. The combined analysis is complementary to the study of individual phenotypes. A combined analysis improves study power by increasing total sample size; on the other hand, study of individual phenotypes may reveal drugs or drug classes that are specifically useful for particular disease symptoms or subtypes, for example melancholic depression.

We performed meta-analysis of *p*-values based on two methods, the Simes’ method ^29^ and the Brown’s approach ^30,31^. The Simes’ method is valid under positive regression dependencies ^32^. Brown’s method is similar to Fisher’s method but also accounts for dependencies in *p*-values. Briefly, assuming *k* p-values, Fisher showed that the statistic *T* = Σ−2 log *P*_*i*_ should follow a chi-squared distribution with *2k* degrees of freedom if the *p*-values are independent. Brown’s method is an extension of Fisher’s approach by estimating the statistic *T* with a re-scaled chi-square distribution 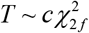 where

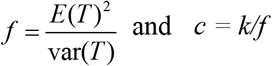

The expectation of the statistic *T* can be estimated by *E*(*T*) = *2k* and the variance by 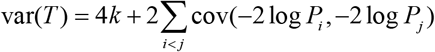. We estimated the covariance empirically from the observed *p*-value vectors of different phenotypes.

The MDD-CONVERGE sample includes only Chinese subjects and does not overlap with other datasets, but we accounted for overlapping samples (with Brown’s or Simes’ method) in the remaining GWAS studies. We did not include MDD-PGC when combining *p*-values as this sample is already included in the GWAS meta-analysis of depressive symptoms ^16^.

### Testing for enrichment of psychiatric and other drug classes

We considered three sources when defining psychiatric drug-sets in our analyses. The first set came from drugs listed in the Anatomical Therapeutic Classification (ATC) system downloaded from KEGG. We extracted three groups of drugs: (1) all psychiatric drugs (coded “N05” or “N06”); (2) antipsychotics (coded “N05A”); (3) antidepressants and anxiolytics (coded “N05B” or “N06A”). The second source was from MEDication Indication resource (MEDI) ^33^ derived from four public medication resources, namely RxNorm, Side Effect Resource 2 (SIDER2), Wikipedia and MedlinePlus. A random subset of the extracted indications was checked by physicians. The MEDI high-precision subset (MEDI-HPS), with an estimated curation precision of 92%, was used in our analyses ^33^. Since only known drug indications are included in ATC or MEDI-HPS, we also included an expanded set of drugs that are considered for clinical trials (as listed on https://clinicaltrials.gov). These drugs are usually promising candidates supported by preclinical or clinical studies. A precompiled list of these drugs was obtained from https://doi.org/10.15363/thinklab.d212. We also examined enrichment for closely related disorders in combinations, including schizophrenia with bipolar disorder (BD), as well depression with anxiety.

The above represents a hypothesis-driven analysis of psychiatric drug classes. To explore whether *other* drug groups may be repositioned for disease treatment, we also performed a comprehensive enrichment analysis of all ATC level 3 drug classes. To avoid the results being driven by too few drugs in a class, we limited the analyses to drug classes with at least 5 members. Note that in the previous step we have found *individual drugs* as repositioning candidates, but we also hope to find out which drug *classes* may be promising for repositioning, which can also provide insights into potentially new mechanisms of actions in future development.

We performed enrichment tests of repositioning hits for known drug classes, in a manner similar to the GSA described above. P-values are first converted to *z*-statistics, and the mean z-score within each drug class is compared against the theoretical null of zero (self-contained test) and against other drugs outside the designated drug class (competitive test) with one-sided tests.

It is reasonable to believe that the current antidepressants or anxiolytics are not the only drugs that have therapeutic effects; in other words, a certain proportion of drugs in the “competing set” might also have therapeutic potential against depression or anxiety. Therefore, results of the competitive tests should be interpreted with this potential limitation in mind. In this paper we presented the drug-set enrichment results of both self-contained and competitive tests.

### Literature search and curation of results

We extracted the top 20 repositioning hits of each psychiatric trait and meta-analyzed results, with drug-related gene-sets derived from either the entire database or drug targets defined by PubChem or ChEMBL. We performed a systematic search in PubMed and Google scholar using the following terms: Drug_name AND (depression OR depressive OR antidepressant OR anxiety OR panic OR phobia OR anxiolytic). References therein were looked up as necessary. The search was performed in June to August 2017.

### Correction for multiple testing

We employed the false discovery rate (FDR) approach (which controls the expected proportion of false positives among those declared to be significant) to account for multiple testing ^34^ FDR-adjusted *p*-values (or *q*-values) were computed by the R function p.adjust with the Benjamini-Hochberg (BH) procedure ^34^. The primary *q*-value threshold was set at 0.05, while results with *q* < 0.1 were regarded as suggestive associations.

## RESULTS

### Enrichment of psychiatric drug classes among the drugs repositioned from gene-set analyses

Table 1–Table 3 and Supplementary Table 1–Table 3 show the enrichment *p*-values and *q*-values for major psychiatric drug classes amongst the drugs repositioned from GSA. We observed that the drugs repositioned from most GWAS of anxiety and depressive traits are generally enriched for known psychiatric medications.

**Table 1.** Enrichment *p*-values of repositioning hits derived from GWAS of major depressive disorder (MDD) and depressive symptoms

| Disorder | MDD-CON | MDD-CON | MDD-PGC | MDD-PGC | DepSym | DepSym |
| --- | --- | --- | --- | --- | --- | --- |
|  | Self | Compet | Self | Compet | Self | Compet |
| *ATC classification* |  |  |  |  |  |  |
| Antipsychotics | **2.50E-04** | **1.13E-02** | 1.00E+00 | 9.98E-01 | **1.35E-03** | *4.48E-02* |
| Antidepressants or anxiolytics | **2.10E-06** | **7.64E-04** | 1.00E+00 | 9.82E-01 | **1.90E-02** | 3.44E-01 |
| All ATC psychiatric drugs | **3.54E-08** | **1.83E-03** | 1.00E+00 | 9.98E-01 | **5.17E-07** | **9.62E-03** |
| *MEDI-HPS* |  |  |  |  |  |  |
| Schizophrenia and Bipolar | **3.06E-07** | **2.04E-04** | 1.00E+00 | 1.00E+00 | **2.90E-02** | 4.68E-01 |
| Anxiety and Depression | **2.98E-07** | **6.59E-04** | 1.00E+00 | 9.99E-01 | **2.08E-03** | 2.02E-01 |
| All psychiatric drugs | **1.77E-10** | **4.40E-05** | 1.00E+00 | 1.00E+00 | **1.35E-03** | 3.15E-01 |
| *ClinicalTrial.gov* |  |  |  |  |  |  |
| Anxiety disorders | **3.20E-07** | **2.18E-03** | 1.00E+00 | 9.92E-01 | **4.97E-04** | 1.16E-01 |
| Depression | **9.45E-07** | **2.27E-02** | 1.00E+00 | 9.99E-01 | **5.76E-05** | 8.67E-02 |
| Bipolar disorder | **1.64E-03** | 1.69E-01 | 1.00E+00 | 9.97E-01 | **1.55E-05** | **9.33E-03** |
| Schizophrenia | **5.42E-05** | 9.66E-02 | 1.00E+00 | 9.96E-01 | **4.39E-05** | 9.42E-02 |
| Anxiety + Depression | **3.14E-09** | **1.69E-03** | 1.00E+00 | 1.00E+00 | **1.23E-05** | *6.07E-02* |
| Schizophrenia + Bipolar | **2.62E-05** | 9.12E-02 | 1.00E+00 | 9.96E-01 | **4.85E-07** | **8.78E-03** |
| All psychiatric drugs | **6.89E-11** | **1.43E-03** | 1.00E+00 | 1.00E+00 | **5.09E-09** | **1.49E-02** |

**Table 3.** Enrichment p-values of repositioning hits derived from meta-analysis of GWAS *p*-values from MDD-CONVERGE, MDD-PGC, SSGAC-DS and SSGAC-NEU

| Disorder | Brown Self | Brown Compet | Simes Self | Simes Compet |
| --- | --- | --- | --- | --- |
| *ATC classification* |  |  |  |  |
| Antipsychotics | **2.31E-09** | **8.77E-07** | **4.64E-07** | **9.11E-06** |
| Antidepressants or anxiolytics | **8.30E-10** | **3.19E-06** | **2.01E-07** | **9.57E-06** |
| All ATC psychiatric drugs | **2.51E-18** | **3.16E-10** | **8.38E-12** | **3.03E-08** |
| *MEDI-HPS* |  |  |  |  |
| Schizophrenia and Bipolar | **5.19E-08** | **5.70E-05** | **7.14E-06** | **1.79E-04** |
| Anxiety and Depression | **1.63E-10** | **3.18E-06** | **1.35E-06** | **8.95E-05** |
| All psychiatric drugs | **2.36E-13** | **2.85E-07** | **1.83E-08** | **6.63E-06** |
| *ClinicalTrial.gov* |  |  |  |  |
| Anxiety disorders | **3.47E-08** | **4.63E-04** | **1.98E-04** | **7.49E-03** |
| Depression | **4.33E-10** | **3.82E-04** | **1.04E-06** | **6.14E-04** |
| Bipolar disorder | **5.53E-09** | **7.43E-05** | **8.07E-07** | **7.47E-05** |
| Schizophrenia | **1.61E-10** | **9.69E-05** | **2.04E-06** | **4.70E-04** |
| Anxiety + Depression | **4.94E-12** | **4.19E-05** | **5.33E-08** | **7.93E-05** |
| Schizophrenia + Bipolar | **2.56E-12** | **6.00E-06** | **5.84E-08** | **3.07E-05** |
| All psychiatric drugs | **1.23E-16** | **9.29E-07** | **2.94E-10** | **4.86E-06** |

First we consider the three datasets (MDD-CONVERGE, MDD-PGC, SSGAC-DS) which focus on depression traits (Table 1 and Supplementary Table 1). On the whole the MDD-CONVERGE sample showed the strongest enrichment with the greatest number of significant results. Significant enrichment was seen for antipsychotics and antidepressants or anxiolytics within ATC and MEDI-HPS categories. In contrast, we did not observe any significant enrichment for drugs repositioned from the MDD-PGC sample. The SSGAC-DS study included MDD-PGC data but the latter only comprised ~10% of the total sample size. For SSGAC-DS, we observed enrichment of drugs for schizophrenia and BD, and suggestive associations with medications for anxiety and depression listed in clinicalTrial.gov.

As for GWAS studies on neuroticism (SSGAC-NEU) and anxiety disorders, there was evidence of enrichment for most psychiatric drug classes under study. Interestingly, for neuroticism, the strongest enrichment was for antipsychotics (lowest *q* = 2.28E-09) instead of antidepressants. Table 2 and Supplementary Table 2 show the enrichment *p*-values and *q*-values respectively.

**Table 2.** Enrichment *p*-values of repositioning hits derived from GWAS of anxiety disorders and neuroticism

| Disorder | AnxietyCC Self | AnxietyCC Compet | Neurotic Self | Neurotic Compet |
| --- | --- | --- | --- | --- |
| *ATC classification* |  |  |  |  |
| Antipsychotics | **1.71E-03** | **4.47E-04** | **1.21E-10** | **1.52E-10** |
| Antidepressants or anxiolytics | **1.35E-02** | **5.23E-03** | **1.09E-03** | **1.45E-03** |
| All ATC psychiatric drugs | **6.67E-03** | **7.93E-04** | **4.88E-12** | **7.99E-12** |
| *MEDI-HPS* |  |  |  |  |
| Schizophrenia and Bipolar | **7.51E-03** | **2.30E-03** | **1.29E-06** | **1.98E-06** |
| Anxiety and Depression | **8.24E-03** | **2.46E-03** | **5.98E-05** | **9.01E-05** |
| All psychiatric drugs | **4.18E-04** | **3.82E-05** | **1.02E-07** | **1.77E-07** |
| *ClinicalTrial.gov* |  |  |  |  |
| Anxiety disorders | **2.84E-02** | **7.94E-03** | **2.70E-03** | **3.82E-03** |
| Depression | **1.17E-02** | **1.55E-03** | 8.62E-02 | 1.14E-01 |
| Bipolar disorder | **1.56E-02** | **3.53E-03** | **5.93E-03** | **7.94E-03** |
| Schizophrenia | **3.49E-03** | **3.43E-04** | **9.70E-04** | **1.47E-03** |
| Anxiety + Depression | **1.78E-02** | **2.33E-03** | *4.09E-02* | *5.70E-02* |
| Schizophrenia + Bipolar | **3.12E-03** | **2.69E-04** | **5.32E-04** | **7.84E-04** |
| All psychiatric drugs | **8.26E-03** | **3.61E-04** | **2.16E-03** | **3.34E-03** |

For analyses involving meta-analyzed GWAS data across all datasets (Table 3 and Supplementary Table 3), enrichment was observed for all psychiatric drug classes, with generally stronger or at least comparable statistical associations when compared to enrichment tests of individual GWAS. The results of Brown’s and Simes’ tests were largely consistent with each other.

We also performed an analysis based on drug target genes derived from PubChem or ChEBML only. The results (including corresponding q-values) were given in Supplementary Table 4 and were largely similar to the above findings.

**Table 4.** Top 5 enriched ATC drug classes with gene-sets derived from the whole DSigDB (considering all drug classes)

| level3_codes | level3_name | pval self | qval self | pval<br>compet | qval<br>compet |
| --- | --- | --- | --- | --- | --- |
| **MDD-CONVERGE** |  |  |  |  |  |
| D07A | CORTICOSTEROIDS, PLAIN | 3.83E-08 | 4.94E-06 | 1.77E-06 | 2.28E-04 |
| D07X | CORTICOSTEROIDS, OTHER COMBINATIONS | 5.04E-07 | 2.17E-05 | 7.56E-06 | 4.88E-04 |
| N06A | ANTIDEPRESSANTS | 3.35E-07 | 2.16E-05 | 7.39E-05 | 2.57E-03 |
| A07E | INTESTINAL ANTIINFLAMMATORY AGENTS | 2.16E-05 | 4.64E-04 | 9.26E-05 | 2.57E-03 |
| S01C | ANTIINFLAMMATORY AGENTS AND<br>ANTIINFECTIVES IN COMBINATION | 1.65E-05 | 4.26E-04 | 9.96E-05 | 2.57E-03 |
| **MDD-PGC** |  |  |  |  |  |
| G01A | ANTIINFECTIVES AND ANTISEPTICS, EXCL.<br>COMBINATIONS WITH CORTICOSTEROIDS | 2.50E-03 | 3.23E-01 | 1.46E-04 | 1.88E-02 |
| M01A | ANTIINFLAMMATORY AND ANTIRHEUMATIC<br>PRODUCTS, NON-STERIODS | 7.20E-03 | 4.64E-01 | 3.35E-04 | 2.16E-02 |
| L01C | PLANT ALKALOIDS AND OTHER NATURAL<br>PRODUCTS | 1.92E-02 | 8.26E-01 | 4.69E-03 | 2.02E-01 |
| N05C | HYPNOTICS AND SEDATIVES | 1.73E-01 | 1.00E+00 | 1.08E-02 | 3.48E-01 |
| L01X | OTHER ANTINEOPLASTIC AGENTS | 2.63E-01 | 1.00E+00 | 1.35E-02 | 3.48E-01 |
| **Depressive symptoms** |  |  |  |  |  |
| N01B | ANESTHETICS, LOCAL | 1.07E-07 | 1.19E-05 | 9.60E-06 | 9.09E-04 |
| D08A | ANTISEPTICS AND DISINFECTANTS | 1.85E-07 | 1.19E-05 | 1.41E-05 | 9.09E-04 |
| V03A | ALL OTHER THERAPEUTIC PRODUCTS | 3.27E-06 | 1.41E-04 | 1.74E-04 | 7.48E-03 |
| C10A | LIPID MODIFYING AGENTS, PLAIN | 6.81E-04 | 1.94E-02 | 4.05E-03 | 1.31E-01 |
| N04B | DOPAMINERGIC AGENTS | 1.29E-03 | 2.18E-02 | 8.24E-03 | 2.13E-01 |
| **Anxiety disorders** |  |  |  |  |  |
| N06A | ANTIDEPRESSANTS | 6.05E-04 | 6.97E-02 | 2.14E-04 | 2.76E-02 |
| N05A | ANTIPSYCHOTICS | 1.71E-03 | 7.35E-02 | 4.47E-04 | 2.88E-02 |
| L01B | ANTIMETABOLITES | 1.08E-03 | 6.97E-02 | 7.03E-04 | 3.02E-02 |
| C07A | BETA BLOCKING AGENTS | 1.19E-02 | 3.17E-01 | 6.31E-03 | 2.03E-01 |
| N04A | ANTICHOLINERGIC AGENTS | 1.23E-02 | 3.17E-01 | 9.13E-03 | 2.32E-01 |
| **Neuroticism** |  |  |  |  |  |
| N05A | ANTIPSYCHOTICS | 1.21E-10 | 1.56E-08 | 1.52E-10 | 1.96E-08 |
| D01A | ANTIFUNGALS FOR TOPICAL USE | 2.98E-04 | 1.92E-02 | 3.31E-04 | 2.13E-02 |
| G01A | ANTIINFECTIVES AND ANTISEPTICS, EXCL.<br>COMBINATIONS WITH CORTICOSTEROIDS | 5.67E-04 | 2.44E-02 | 6.22E-04 | 2.67E-02 |
| N05C | HYPNOTICS AND SEDATIVES | 3.31E-03 | 1.07E-01 | 4.03E-03 | 1.30E-01 |
| N06A | ANTIDEPRESSANTS | 6.15E-03 | 1.58E-01 | 7.84E-03 | 1.69E-01 |
| **Simes p** |  |  |  |  |  |
| N05A | ANTIPSYCHOTICS | 4.64E-07 | 3.82E-05 | 9.11E-06 | 7.35E-04 |
| N06A | ANTIDEPRESSANTS | 5.93E-07 | 3.82E-05 | 1.14E-05 | 7.35E-04 |
| G01A | ANTIINFECTIVES AND ANTISEPTICS, EXCL.<br>COMBINATIONS WITH CORTICOSTEROIDS | 4.53E-04 | 1.95E-02 | 1.24E-03 | 5.33E-02 |
| A01A | STOMATOLOGICAL PREPARATIONS | 7.37E-03 | 1.64E-01 | 1.74E-02 | 3.43E-01 |
| V03A | ALL OTHER THERAPEUTIC PRODUCTS | 3.94E-03 | 1.27E-01 | 1.93E-02 | 3.43E-01 |
| **Brown's p** |  |  |  |  |  |
| N05A | ANTIPSYCHOTICS | 2.31E-09 | 2.98E-07 | 8.77E-07 | 1.13E-04 |
| N06A | ANTIDEPRESSANTS | 6.41E-09 | 4.13E-07 | 3.58E-06 | 2.31E-04 |
| G01A | ANTIINFECTIVES AND ANTISEPTICS, EXCL.<br>COMBINATIONS WITH CORTICOSTEROIDS | 1.44E-04 | 4.64E-03 | 1.04E-03 | 4.47E-02 |
| V03A | ALL OTHER THERAPEUTIC PRODUCTS | 7.00E-05 | 3.01E-03 | 2.86E-03 | 9.22E-02 |
| N04B | DOPAMINERGIC AGENTS | 1.61E-03 | 3.46E-02 | 9.57E-03 | 2.24E-01 |

Table 4 and Supplementary Table 5 shows the results of enrichment tests across all ATC level 3 drug classes based on gene-sets derived from the whole DSigDB database. Table 5 and Supplementary Table 6 shows the findings from the same analysis but with gene-sets derived from PubChem or ChEBML only. We consider results from both of the analyses here. For MDD-PGC, anti-infective and anti-inflammatory agents were significantly enriched. Interestingly, hypnotics/sedatives and anxiolytics, which were commonly used as short-term therapies especially at the acute phase of illness^35^, are also ranked among the top. Antidepressants or other psychiatric drugs however were not enriched. For MDD-CONVERGE, drug classes pertaining to corticosteroids were ranked highly. Notably, antidepressants and antipsychotics were ranked within the top 5 in our two sets of analyses. For anxiety disorders, antipsychotics and antidepressants were the two most strongly enriched medication classes. Beta-blocking agents were ranked among the top for anxiety disorders. Beta adrenergic blockers, especially propranolol, have been clinically used in anxiety disorders for a long time, although the efficacy still remains uncertain^36^. As for depressive symptoms, dopaminergic and antiepileptic agents were listed among the top, and interestingly lipid-lowering agent was also on the top list. As for neuroticism, antipsychotics was the most strongly enriched drug class, and antidepressants was also ranked within the top 10. Other drug classes listed in top 10 also included hypnotics/sedatives, anxiolytics and dopaminergic agents. The full results are shown in Supplementary Table 5 to 6.

**Table 5.** Top 5 enriched ATC drug classes with gene-sets derived from PubChem and ChEMBL (considering all drug classes)

| level3 | level3_name | pval self | qval self | pval | qval |
| --- | --- | --- | --- | --- | --- |
| codes |  |  |  | compet | compet |
| **MDD-PGC** |  |  |  |  |  |
| N01A | ANESTHETICS, GENERAL | 4.06E-03 | 2.05E-01 | 2.94E-04 | 2.32E-02 |
| N05C | HYPNOTICS AND SEDATIVES | 6.16E-02 | 1.00E+00 | 7.86E-04 | 2.77E-02 |
| L01C | PLANT ALKALOIDS AND OTHER<br>NATURAL PRODUCTS | 5.19E-03 | 2.05E-01 | 1.05E-03 | 2.77E-02 |
| N05B | ANXIOLYTICS | 6.25E-02 | 1.00E+00 | 1.68E-03 | 3.00E-02 |
| L01X | OTHER ANTINEOPLASTIC<br>AGENTS | 6.35E-01 | 1.00E+00 | 1.90E-03 | 3.00E-02 |
| **MDD-CONVERGE** |  |  |  |  |  |
| N05A | ANTIPSYCHOTICS | 5.54E-05 | 4.38E-03 | 1.93E-04 | 1.52E-02 |
| D08A | ANTISEPTICS AND<br>DISINFECTANTS | 2.03E-03 | 7.90E-02 | 3.79E-03 | 1.50E-01 |
| A07E | INTESTINAL<br>ANTIINFLAMMATORY AGENTS | 3.00E-03 | 7.90E-02 | 7.10E-03 | 1.87E-01 |
| A03A | DRUGS FOR FUNCTIONAL<br>GASTROINTESTINAL DISORDERS | 1.25E-02 | 1.98E-01 | 1.51E-02 | 2.98E-01 |
| G03X | OTHER SEX HORMONES AND<br>MODULATORS OF THE GENITAL<br>SYSTEM | 1.19E-02 | 1.98E-01 | 1.99E-02 | 3.14E-01 |
| **Anxiety case-control** |  |  |  |  |  |
| N05A | ANTIPSYCHOTICS | 5.23E-07 | 4.13E-05 | 3.27E-08 | 2.58E-06 |
| N06A | ANTIDEPRESSANTS | 5.24E-04 | 1.53E-02 | 1.17E-04 | 4.62E-03 |
| D01A | ANTIFUNGALS FOR TOPICAL USE | 5.81E-04 | 1.53E-02 | 2.42E-04 | 6.37E-03 |
| J02A | ANTIMYCOTICS FOR SYSTEMIC USE | 3.39E-03 | 6.70E-02 | 2.52E-03 | 4.25E-02 |
| N02C | ANTIMIGRAINE PREPARATIONS | 6.06E-03 | 9.57E-02 | 2.69E-03 | 4.25E-02 |
| **Depressive symptoms** |  |  |  |  |  |
| N04B | DOPAMINERGIC AGENTS | 1.03E-04 | 4.05E-03 | 1.46E-03 | 7.79E-02 |
| N03A | ANTIEPILEPTICS | 1.01E-04 | 4.05E-03 | 2.62E-03 | 7.79E-02 |
| L01B | ANTIMETABOLITES | 3.34E-04 | 5.36E-03 | 2.96E-03 | 7.79E-02 |
| M03A | MUSCLE RELAXANTS, PERIPHERALLY ACTING AGENTS | 2.87E-03 | 2.22E-02 | 4.82E-03 | 9.22E-02 |
| L01A | ALKYLATING AGENTS | 2.08E-03 | 2.05E-02 | 5.83E-03 | 9.22E-02 |
| **Neuroticism** |  |  |  |  |  |
| N05A | ANTIPSYCHOTICS | 2.64E-13 | 2.09E-11 | 4.31E-10 | 3.40E-08 |
| N05B | ANXIOLYTICS | 9.25E-04 | 3.65E-02 | 6.92E-03 | 2.73E-01 |
| N05C | HYPNOTICS AND SEDATIVES | 1.70E-03 | 4.48E-02 | 1.39E-02 | 3.66E-01 |
| N04B | DOPAMINERGIC AGENTS | 7.60E-03 | 1.00E-01 | 2.99E-02 | 5.78E-01 |
| R06A | ANTI HISTAMINES FOR SYSTEMIC USE | 3.38E-03 | 5.40E-02 | 3.66E-02 | 5.78E-01 |
| **Simes p** |  |  |  |  |  |
| N05A | ANTIPSYCHOTICS | 4.98E-08 | 3.93E-06 | 1.95E-06 | 1.54E-04 |
| N04B | DOPAMINERGIC AGENTS | 1.90E-04 | 7.51E-03 | 5.58E-04 | 2.20E-02 |
| L01A | ALKYLATING AGENTS | 1.90E-03 | 4.86E-02 | 3.19E-03 | 8.40E-02 |
| L02B | HORMONE ANTAGONISTS AND RELATED AGENTS | 3.32E-03 | 5.25E-02 | 5.36E-03 | 1.06E-01 |
| N02C | ANTIMIGRAINE PREPARATIONS | 2.46E-03 | 4.86E-02 | 7.81E-03 | 1.23E-01 |
| **Brown p** |  |  |  |  |  |
| N05A | ANTIPSYCHOTICS | 7.68E-11 | 6.07E-09 | 6.57E-09 | 5.19E-07 |
| N04B | DOPAMINERGIC AGENTS | 7.68E-05 | 3.03E-03 | 2.87E-04 | 1.13E-02 |
| L02B | HORMONE ANTAGONISTS AND<br>RELATED AGENTS | 3.14E-03 | 4.96E-02 | 5.99E-03 | 1.44E-01 |
| N02C | ANTIMIGRAINE PREPARATIONS | 1.64E-03 | 3.24E-02 | 7.30E-03 | 1.44E-01 |
| N06A | ANTIDEPRESSANTS | 1.08E-03 | 2.84E-02 | 1.68E-02 | 2.65E-01 |

**Table 6.**
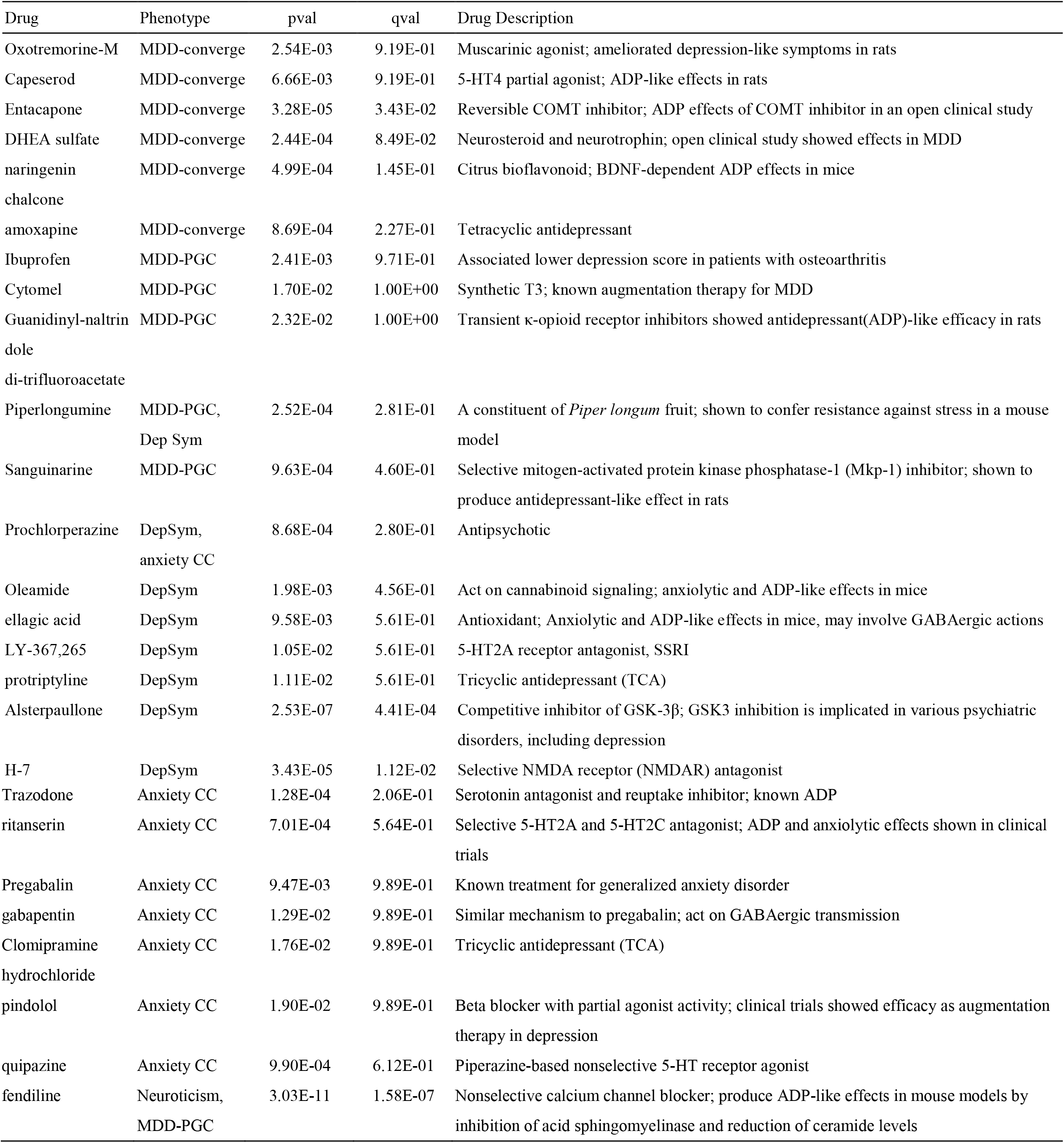
Selected repositioning hits with literature support

## Top repositioning hits

### Based on gene-sets derived from the whole DSigDB database

We found several interesting repositioning hits that were supported by previous studies (Table 6; Supplementary Table 7 gives a fully annotated list of the top 20 drugs). The top-ranked repositioning hit identified in the meta-analysis was fendiline (Brown’s *p* = 1.06E-11, *q* = 5.55E-8), a non-selective calcium channel blocker. Fendiline was shown to exert antidepressant-like effects in a mouse model by inhibition of acid sphingomyelinase (ASM) activity and reduction of ceramide concentrations in the hippocampus ^37^. A drop in ceramide concentrations might lead to increased neurogenesis and improved neuronal maturation and survival ^37^. The drug was ranked among the top for MDD-PGC, SSGAC-DS and neuroticism.

As for individual phenotypes, some interesting candidates for MDD-CONVERGE include DHEA, a neurosteroid with evidence of antidepressive effects in a double-blind RCT; naringenin chalcone, a citrus bioflavonoid shown to be effective in mouse models^38^; amoxapine, a tetracyclic antidepressant^39^.

Candidates for MDD-PGC include ibuprofen, an NSAID shown to be associated with lower depressive symptoms in osteoarthritis patients; piperlongumine, a constituent of the fruit of *Piper longum*, which was shown to confer resistance against stress in a mouse model ^40^; and sanguinarine, a selective mitogen-activated protein kinase phosphatase-1 (Mkp-1) inhibitor which produced antidepressant-like effect in rats^41^.

For anxiety disorders, candidates included a known antidepressant trazodone and a serotonin agonist quipazine that may increase brain serotonin levels^42^. For depressive symptoms, alsterpaullone, a glycogen synthase kinase-3β (GSK-3β) inhibitor ^43^. Increased activation of GSK-3*β* was also associated with depression-like behavior in mouse models, which could be alleviated by GSK-3*β* inhibitors ^44^. In addition, inhibition of GSK3 has been postulated as a major mechanism of action by the mood stabilizer lithium ^45^. A NMDA receptor antagonist H-7 was also top-listed^46^.

Among the top repositioning hits from the individual and meta-analysis results, a number of them are calcium channel blockers (CCB). These include fendiline, perhexiline, prenylamine and felodipine (prenylamine was withdrawn from the market due to risk of QT prolongation and torsades de pointes ^47^). Although with the exception of fendiline, no direct experimental or clinical studies have shown antidepressant or anxiolytic properties of the above drugs, CCB as a whole have been proposed as treatment for various psychiatric disorders. CCB has been mostly studied for the treatment of mania, recently reviewed in Cipriani et al. ^48^. However the number of quality double-blind randomized controlled trials (RCT) was small, and there is yet no sufficient evidence to suggest the use of CCB in treating manic symptoms. As for depression, a recent pilot (patient-only) study of isradipine on bipolar depression showed positive results ^49^. Another CCB, nicardipine, was reported to enhance the antidepressant action of electroconvulsive therapy^50^. Notwithstanding the mixed evidence, CCB are probably still worthy of further investigation for depression and anxiety disorders, given the biological relevance of calcium signaling, preliminary support from clinical studies and the wide availability as well as known safety profiles of this drug class.

### Based on Gene-sets derived from PubChem and ChEMBL

A manually annotated list of the top 20 prioritized drugs is given in supplementary Table 8. Some highlighted candidates are also shown in Table 6. Some repositioning hits for MDD-CONVERGE included oxotremorine-M, a muscarinic acetylcholine receptor agonist that may ameliorate depressive symptoms and restore hippocampal neurogenesis in an animal model^51^ and capeserod, a 5-HT4 receptor agonist shown effective in a mouse model^52^. For MDD-PGC, top-listed drugs include three NSAIDs, namely ibuprofen, meclofenamate and flufenamic acid, which possess anti-inflammatory actions that may also be beneficial for depression^53,54^. Remarkably, our analysis recovered a known augmentation therapy for depression, namely Cytomel (or synthetic triiodothyronine [T3])^55^. Another candidate was guanidinonaltrindole di-trifluoroacetate, a κ-opioid receptor inhibitor with antidepressant-like and anxiolytic-like efficacy in rat models^56^.

There were a number of noteworthy candidates from the analysis results of anxiety disorders. Notably, the top-ranked drugs contain pregabalin and gabapentin, two drugs with similar mechanisms which have been shown anxiolytic effects in clinical trials (especially pregabalin)^57,58^; clomipramine, a known tricyclic antidepressant; and trazodone, another known antidepressant belonging to the serotonin antagonist and reuptake inhibitor class. Some other noteworthy candidates include ritanserin, a selective 5-HT_2A_ and 5-HT_2C_ receptor antagonist^59–61^; pindolol, a beta blocker with partial beta-adrenergic receptor agonist activity and a possible augmentation therapy for depression^62,63^; atomoxetine, a norepinephrine reuptake inhibitor.

For depressive symptoms, the top-listed candidates include prochlorperazine and raclopride, which are antipsychotics; protriptyline, a tricyclic antidepressant; oleamide, a cannabinoid receptor type 1 (CB1) receptor agonist of potential antidepressant effects in animal models^64,65^; ellagic acid and chebulinic acid (an ellagitannin), natural phenol antioxidants with some evidence of improving depressive traits again in animal models^66–68^. Also of note is the nicotinic agonist DMPP, as nicotine has been shown to have antidepressant properties in pre-clinical and clinical studies^69^; the antidepressant effects may be due to initial activation of the nicotinic receptor followed by densensitization leading to long term antagonism^69^. For neuroticism, again quite a number of top hits are CCB, and the CCB fendiline was the top-ranked candidate. The repositioning hits derived from meta-analyzed *p*-values were covered above.

## DISCUSSION

In this study we leveraged large-scale GWAS summary data and analyzed gene-sets associated with drugs to uncover repositioning opportunities for depression and anxiety disorders. It is encouraging that we observed significant enrichment for known psychiatric medications or drugs considered in clinical trials. Remarkably, antipsychotics and antidepressants were the two most significantly enriched drug classes in our combined analysis (with gene-sets derived from the whole DSigDB). Our findings provide support for the validity of GSA in drug repurposing. In addition, we reveal a number of interesting candidates for repurposing that are supported by prior studies. Although relatively few susceptibility variants of genome-wide significance have been found for depression and anxiety disorders, our findings suggest that leveraging variants with weaker associations, for example by GSA, might still contribute valuable information to the discovery of novel therapies.

While we have included three datasets (SSGAC-DS, MDD-PGC, MDD-CONVERGE) directly related to depression, the enrichment results are quite different. Both MDD-PGC and MDD-CONVERGE are case-control GWAS studies on MDD; however, the drugs and drug classes enriched were quite different. The enrichment for psychiatric drug classes were generally weaker for MDD-PGC. The discrepancy might be due to the differences between the two samples. As described above, the two samples differ by gender, ethnicity and the severity of depression. In addition, due to the lower awareness and possibly stronger resistance to seeking medication attention for depression in China, the disease severity in the CONVERGE cohort may be even higher than expected. It is widely accepted that MDD is a heterogeneous disorder, with a variety of clinical presentations and possibly divergent pathophysiologies ^70^. By recruiting a more homogeneous group of patients, the CONVERGE study might have better power in detecting susceptibility genes despite a lower sample size. Indeed, MDD-CONVERGE revealed two genome-wide significant loci while none was found in the MDD-PGC study. It is also worth mentioning that previous meta-analyses showed that the response to antidepressant depends on the baseline severity of depression ^71,72^. The studies reported that effects of antidepressants were largest for the most severely depressed group, but smaller for mild to moderate depression. Our findings, although based on a different study paradigm, are broadly in line with this clinical observation.

It is noteworthy that the repositioning hits are not only enriched for antidepressants or anxiolytics but also antipsychotics. A meta-analysis by Spielmans et al. revealed that atypical antipsychotics are effective as adjunctive treatment for treatment-resistant depression^73^. Zhou et al. also reached a similar conclusion in a recent network meta-analysis^74^. Atypical antipsychotics may also be useful for anxiety disorders and symptoms ^75–77^, although further studies are required and that the benefits need to be balanced against the side-effects. Furthermore, a shared genetic basis between schizophrenia and depression is well-established ^78^, and a recent study also found significant genetic correlation between neuroticism and schizophrenia ^79^. Epidemiology studies also demonstrated associations of neuroticism with schizophrenia^80^.

Several other top-ranked drug classes are also worth mentioning. For MDD-PGC, although antidepressant was not significantly enriched, anti-inflammatory agents was listed among the top. Numerous studies have suggested a role of inflammation in depression, which were reviewed elsewhere^81^. For example, blood levels of inflammatory biomarkers (e.g. IL-1beta, IL-6, TNF, C-reactive protein) were elevated in patients with depression^81^. A large-scale observational study suggested that non-steroidal inflammatory drugs (NSAID), particularly low-dose acetylsalicylic acid, may reduce risk of depression^82^. A meta-analysis of RCTs also reported beneficial effects of anti-inflammatory treatment including both NSAID and cytokine inhibitors^83^.

Another interesting finding is that drug classes related to corticosteroids were highly ranked for MDD-CONVERGE, which is composed of mainly severe melancholic depressive patients. A number of studies^84–86^ have shown that hyperactivity of the hypothalamic-pituitary axis is an important feature of melancholic depression. As with other kinds of gene-set analyses, the current approach of GSA does not delineate the exact direction of drug effects. It may be concluded that corticosteroids or drugs targeting relevant pathways might be of clinical significance to melancholic depression; however, whether corticosteroids themselves or drugs blocking their actions would be useful remains to be investigated. Glucocorticoid synthesis inhibitors and receptor antagonists have been shown to exhibit antidepressant effects^87,88^. Conversely, several short-term trials or studies reported antidepressant properties of corticosteroids themselves; one possible mechanism is the restoration of negative feedback of the HPA axis at the hypothalamic and pituitary levels^89–92^. In any case, our findings provide a proof-of-concept example that making use of appropriate GWAS (or other human genomic) data might help to develop more targeted treatment for disease subtypes.

A few other drug classes may also be worthy of attention. Dopaminergic agents were ranked among the top for depressive symptoms; stimulants including dopaminergic agents have been tested in clinical trials for unipolar and bipolar depression^93–96^. We also observed that anti-migraine medications were highly ranked in the list. An increased rate of depression among migraine patients is well-established^97^, and antidepressants (mainly tricyclics) have been used for migraine prophylaxis^98^. The effects of anti-migraine medications on depressive symptoms however remains to be elucidated. Lipid-lowering agents was top-listed for depressive symptoms (based on all gens in DSigDB) and anxiety disorders (based on drug target genes from PubChem and ChEMBL), and studies have shown potential benefits of statins in depression^99^, and when used with concomitant SSRI^100^. However, some studies have also reported increased depressive symptoms with statins^101^, hence the exact relationship may be complex and may differ by patient characteristics. Whether other types of lipid-lowering drugs may be beneficial is another topic worthy of further investigations.

In this study we employed the GSA approach to drug repositioning. The current study is complementary to our recent repositioning attempt using a new framework in which the drug-induced transcriptome is compared against GWAS-imputed expression profiles^102^. Each of these two methods has their own advantages and disadvantages. The methodology of finding reversed expression patterns (as detailed in So et al.^102^) has a unique advantage of accounting for the directions of associations. It also takes into account the functional impact of variants on expression and is intuitive from a biological point of view. While differentially expressed genes can be included in gene-sets, the actual (quantitative) expression changes are not considered which results in a loss of information. GSA also does not delineate the directions of effects. Nevertheless, GSA can directly make use of knowledge concerning known drug targets and other information on drug-related genes, for which more databases are available. Moreover, in the ‘reversed transcriptome’ approach^102^, one only considers functional effects of the SNPs *on expression*, but genetic variants may also contribute to disease pathogenesis via other mechanisms, such as splicing or changes in protein function. The gene-based test in the current analysis is based on combining statistical evidence of individual SNPs in a general manner, and may capture a wider range of effects on disease risk. Also, the transcriptome comparison approach involves “imputing” expression levels; since the major reference transcriptome dataset (GTEx) is mainly composed of Caucasians (84.6%) with greater proportion of males (65.6%) (https://www.gtexportal.org/home/tissueSummaryPage, accessed 7^th^ Sep 2017), the quality of imputation for other ethnicities and females might be less reliable, for example when applied to the MDD-CONVERGE dataset.

Just as medications acting on different pathways might have synergistic therapeutic effects, we believe that it is beneficial to have different approaches for computational drug repositioning to complement each other. Of course, computational methods leveraging human genomic data are not the only means to drug discovery. We believe that a combination of a variety of approaches, including experimental and computational ones, is required to speed up drug repurposing and discoveries.

Our enrichment analyses support the application of GSA in drug repositioning in depression and anxiety. However, we stress that our repositioning results should be validated in further pre-clinical and clinical studies before translation to practice. GSA analyses do not provide information on the direction of effects, as discussed previously. Measures of statistical significance also do not provide definitive evidence for the actual therapeutic effects of the repositioned drugs.

In summary, we have performed a drug repositioning analyses on depression and anxiety disorders, using a gene-set analysis approach considering five related GWAS studies. We showed that the repositioned drugs are in general enriched for known psychiatric medications or those considered in clinical trials. Remarkably, antipsychotics and antidepressants were ranked among the top even if we considered all level 3 ATC drug classes. The results lend further support to the usefulness of human genomic data in guiding drug development in psychiatry, and we hope that the rapid advances in psychiatric genomics research will translate into benefits for patients in the foreseeable future.

## Acknowledgements

This work is partially supported by the Lo-Kwee Seong Biomedical Research Fund and a Direct Grant from the Chinese University of Hong Kong. We also thank Professor Stephen K.W. Tsui and the Hong Kong Bioinformatics Centre for computing support. We would also like to acknowledge the Psychiatric Genomics Consortium, the CONVERGE Consortium, the Social Science Genetics Association Consortium and Otawa et al. for providing open access to full GWAS summary results.

## Author contributions

H.-C.S. conceived and designed the study. H.-C.S. performed data analyses with assistance from C.K.L.C. H.-C.S. interpreted the data. A.L. and S-Y.W. performed drug annotations. H.-C.S. wrote the manuscript and supervised the study.

